# Artificial Intelligence for automatic movement recognition: a network-based approach

**DOI:** 10.1101/2025.02.27.640538

**Authors:** Emahnuel Troisi Lopez, Roberta Minino, Mario De Luca, Francesco Tafuri, Giuseppe Sorrentino, Pierpaolo Sorrentino, Marie-Constance Corsi

**Affiliations:** Department of Education and Sport Sciences, Pegaso Telematic University, 80143, Naples, Italy; Department of Medical, Motor and Wellness Sciences, University of Naples “Parthenope”, 80133 Napoli, Italy; Heracle Lab Research in Educational Neuroscience, Niccolò Cusano University, 00166 Rome, Italy; Department of Economics, Law, Cybersecurity and Sports Sciences, University of Naples “Parthenope”, 80035 Nola, Italy; Institute of Applied Sciences and Intelligent Systems, National Research Council, 80078 Pozzuoli, Italy; ICS Maugeri Hermitage Napoli, via Miano, 80145 Naples, Italy; Aix Marseille University, INSERM, INS, Institut de Neurosciences des Systèmes, 13007, Marseille, France; Department of Biomedical Sciences, University of Sassari, 07100, Sassari, Italy; Sorbonne Université, Institut du Cerveau—Paris Brain Institute—ICM, CNRS, Inria, Inserm, AP-HP, Hôpital de la Pitié Salpêtrière, F-75013 Paris, France

**Keywords:** human movement recognition, network theory, artificial intelligence, movement classification

## Abstract

**Introduction:** automatic movement recognition is often used to support various fields such as clinical, sports, and security. To date, there is a lack of a classification feature that is both interpretable and not movement-specific, characteristics that would enhance generalization and adaptability. Previous studies on motion analysis have shown that coordination properties extracted from full-body movement using network theory can describe specific movement characteristics, making coordination a potential feature for classification.

**Methods:** therefore, we leveraged kinematic data from 168 individuals performing 30 different movements, published in an online dataset. Using network theory, we reduced data dimensionality, obtaining a coordination matrix called the kinectome. By applying support vector machine algorithms, we compared the classification performance of the kinectome with that of principal component analysis, used as an alternative data reduction method.

**Results:** the classification accuracy of the kinectome (0.99 ± 0.01) was significantly higher (pFDR < 0.001) than that of PCA (0.96 ± 0.04). Moreover, unlike PCA, the kinectome demonstrated resilience to data loss, robustness to derived measures, independence from the classification algorithm, and clear interpretability of features.

**Discussion:** our results suggest that kinnectome-based features could capture interpretable changes between movements that could pave the way to new automatic movement recognition approaches dedicated to a wide range of applications, in particular sport training and physical readaptation, and designed for non-data scientists experts.

## 1. Introduction

Movement recognition refers to identifying and interpreting human movements or gestures using sensor data or video inputs, often via machine learning algorithms. The recognition of patterns in motion analysis holds significant utility across a wide range of applications, including ergonomics, security, process automation, sport, and clinical evaluation ^1^. An effective recognition of motion patterns provides insights into the characteristics of the movements and enables the detection of significant changes from expected patterns. Moreover, the automatic recognition of correct patterns facilitates the assessment of movement quality, efficiency, and effectiveness, serving as targeted feedback based on the type of movement ^2^. Such assessments are crucial in functional reeducation in a hospital setting. Identifying abnormal movement patterns is fundamental to prevent injury, or recognize pathophysiological processes ^3,4^. However, differentiating highly similar patterns is a challenging task that typically requires a substantial amount of data. Generally, the information used to discriminate motion patterns relies on kinematic data, which describes the movement of different body parts in space. Such data can be easily obtained from standard video cameras, which are widely available. However, managing this data may be challenging given the large number of degrees of freedom of the human joints and the synergies needed to execute specific motor tasks ^5^. Indeed, every movement is prescribed by the structure of different body parts, which collaborate to accomplish the intended motion ^6^.

To date, when performing classification of movements, it is common to extract and combine different synthetic features obtained from kinetic and/or kinematic time series. Commonly employed features include peaks of the joint excursion, spatiotemporal characteristics of movement (e.g., gait parameters such as step length), maximum or mean force, speed, or acceleration of specific body parts ^7^. To improve performance, many parameters might be combined, or the entire time series utilized. In both cases, dimensionality reduction is needed for the identification to converge and in order to speed up the process. Indeed, such data often involves tens/hundreds of thousands of values for just a few seconds or minutes of recording ^8^. Selecting the most informative features and reducing dimensionality, while retaining relevant information, poses relevant challenges. One of the main approaches involves principal component analysis (PCA) ^9^. Such a technique reduces the amount of data, creating a new set of variables (i.e., components), of which a few are expected to explain most of the variance of the original data. The resulting components are then used to feed the classification algorithm. However, the PCA components are not directly interpretable, as they represent combinations of the original features ^10^. Hence, further steps must be taken to identify the most relevant features to be used for movement recognition and, consequently, to determine the best methodological approaches for their extraction.

The choice of the features might vary as a function of arbitrary choices, the objectives, and the movements taken into account (that can highly vary among studies). These aspects make the studies difficult to replicate and interpret. Therefore, there is a need to identify alternative features that can be extracted regardless of the type of movement (i.e., they do not depend on specific movements). Furthermore, such a feature should be interpretable and, ideally, intuitively visualizable. This would facilitate explainability, even to clinicians or evaluators that are not specialized in data science. In a previous paper, we showed that it was possible to characterize movement utilizing tools borrowed from network theory ^11^, namely creating a matrix of movement coordination, based on Pearson correlation coefficients (PCC) among pairs of time series representing the kinematics of several body parts ^12^. These matrices, named kinectomes, allowed an almost perfect (up to 99.8%) success rate in differentiating individuals performing a simple walking task. Furthermore, the kinectomes proved to be informative of the executed movement (i.e., gait) and provided a readable and precise information concerning the specific body parts and the pairwise coordination. Hence, by analyzing the network structure, it becomes possible to identify recurring patterns or motifs that are specific to different movements.

Here, we aim to explore the performance of features extracted utilizing the kinectome for pattern recognition of human movement. Building on the proven interpretability of the kinectome, as demonstrated in several previous studies ^12–14^, we set out to test the following hypothesis: i) the classification based on kinectomes would outperform the classical one based on PCA components regardless the classifier; ii) kinectomes would offer a clear visualization of the features, facilitating the interpretability of the results; iii) with respect to PCA components, our approach would demonstrate a better robustness towards missing data and the diversity of the features;. To test our hypothesis, we employed a publicly available dataset including 168 participants performing 30 different movements ^15^, captured by a stereophotogrammetric system. By applying network theory, we constructed a subject-specific movement matrix for each task, based on the time series of the joint excursion.

## 2. Methods

### 2.1 Dataset

In this study, we utilized a dataset published by Zhao et al. ^15^. Briefly, the dataset comprises kinematic time series recorded using a stereophotogrammetric system, capturing 30 different movements performed by 183 athletes (Figure 1). The movements include both mobility and stability tasks, such as back bends, shoulder abductions, and drop jumps. Detailed information about the acquisition system, the sample population, and the specific movements performed can be found in the original paper ^15^. Here, we focused on the time series of joint excursion angles for twelve specific joints: the neck, trunk, and bilateral shoulders, elbows, hips, knees, and ankles. For each joint, three-dimensional time series were available (i.e., anteroposterior, mediolateral, and vertical).

**Figure 1.**
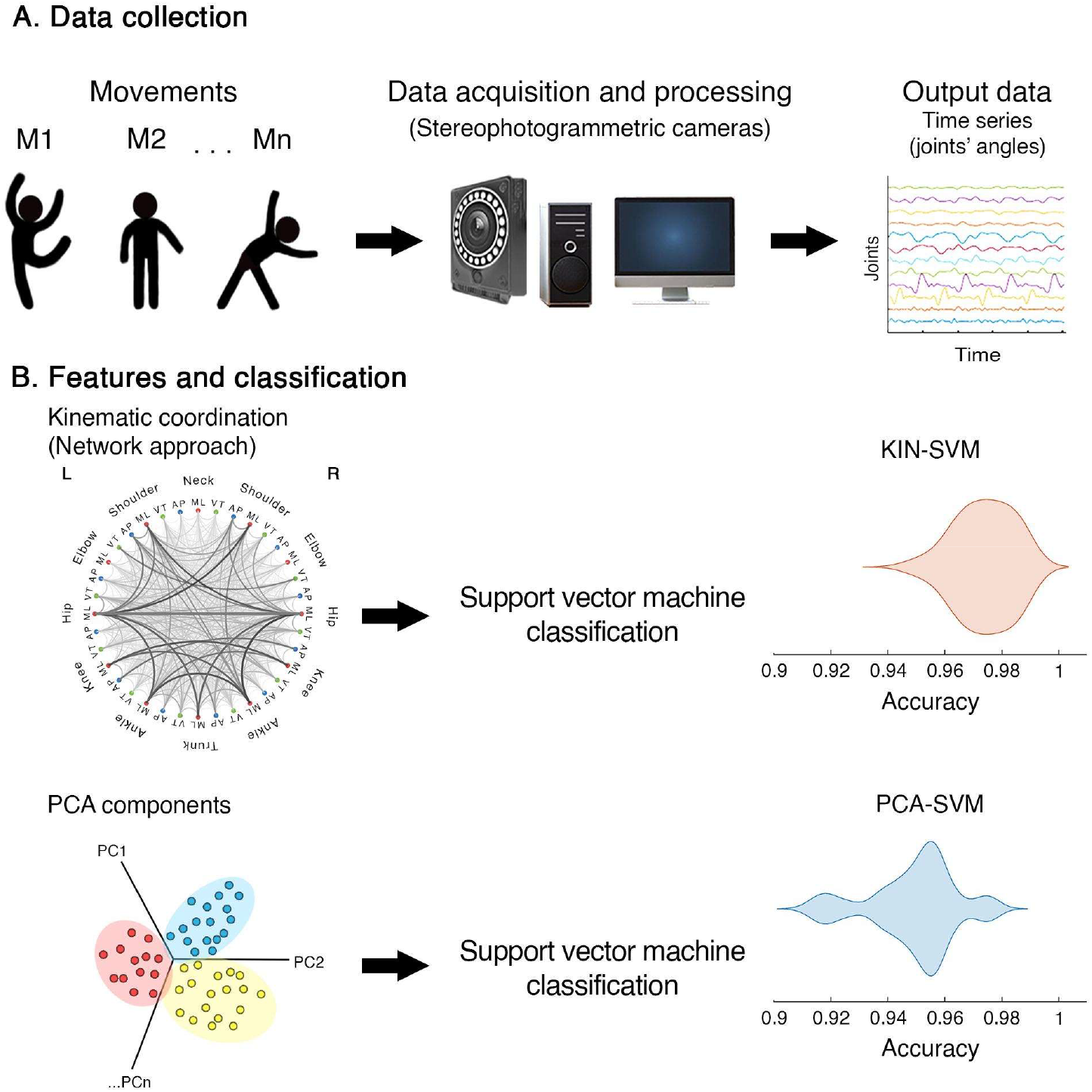
Analysis pipeline. (A) describes data collection related to the online dataset including 30 movements performed by 183 athletes ^15^. After a quality check, the kinematic time series of joint excursions from 168 participants were included in the analysis. (B) delineates the selected feature extraction and classification methodologies. Time series dimensionality was reduced using two different approaches, resulting in two features: edges from the kinematic network representing movement coordination, and principal components extracted through principal component analysis, retaining up to 95% of the variance of the original data. Finally, we determined which feature achieved the highest classification accuracy in movement classification using a support vector machine (SVM).

The data were individually inspected to check for label mismatches, missing values, abnormalities, and overall consistency (e.g., ensuring consistent ordering of data). Missing values were interpolated using MATLAB’s ‘fillmissing’ function with the ‘spline’ method if the gap was less than 0.1 seconds; otherwise, the participant’s data were discarded. After this screening process, 15 individuals were excluded, resulting in a final sample of 168 participants, performing 30 different movements (one recording per movement per participant). As the final step of data preparation, considering that PCA requires data of the same length, we equalized the length of all time series based on the shortest duration (i.e., 68 data points) using MATLAB’s ‘interp1’ function. Various features were then extracted from this dataset for classification analysis.

### 2.2 Features extraction

The first feature we extracted is based on the concept of motor coordination. Drawing from previous works that have employed network theory to measure motor coordination ^12,13,13,14^, we built a kinematic network using the time series of the joints’ angles on the three axes. Joints constitute the node of the network, while their pair-wise coordination is represented by the network’s edges. For each movement, the edges are obtained by calculating the Pearson correlation coefficient between each pair of time series (i.e., 36 in total, given by 12 joints on 3 axes). With this procedure, we built a matrix (i.e., kinectome) for each movement of each participant. Given the symmetrical nature of the matrix, we only considered its upper triangular part for the analysis. These data were then used for movement classification.

To assess the extent to which the kinectome could be considered as a relevant feature for movement recognition, we compared its classification performance with the one resulting from state-of-the-art features, namely, the PCA. We applied the PCA on time series, and we considered the components as features. Specifically, we reorganized the data so that the rows represent the participants and movements (168 participants x 30 movements = 5040 rows), while the columns represent the data points of all time series for each joint (68 data points x 12 joints x 3 dimensions = 2448 columns). Using PCA, we calculated the component scores, which were then used as features. We chose to use a number of components that would explain up to 95% of the variance in the original data. It is important to note that during classification, PCA was applied separately to the training and test sets to avoid overfitting. Additionally, the classification analysis was repeated by recalculating both features on the first (velocity) and second (acceleration) derivatives of the original time series.

### 2.3 Classification

The classification step was performed using a linear Support Vector Machine (SVM) classifier ^16,17^. The classification performances were obtained by a cross-validation that consisted of a stratified k-fold. Such an approach ensures an approximately equivalent of samples of each target (here, of each movement) as the whole dataset. Samples of each class were shuffled before splitting into batches. To ensure the reproducibility of our findings, we set the randomness of the splitting with a fixed pseudo random number generator. We performed this operation 50 times (corresponding to the number of splits).

The performances of the classification were assessed via the accuracy classification score, and the Area under the Receiver Operating Characteristic Curve (ROC AUC) from the prediction score.

To examine the interpretability of the classification performance, we investigated the relative importance of the features, obtained from the absolute value of the classification coefficients of the classification model. We calculated the median value of the feature importance across the cross-validation splits of the testing set.

### 2.4 Benchmark

The main objective of this study is to investigate the discriminant power of two types of features, based respectively on time series and on kinectomes, in the context of movement recognition. Therefore, we worked with one of the simplest classification tools, namely SVM. Nevertheless, to ensure that the obtained classification performances were not dependent on the chosen classifier, we led a benchmark study with the following pipelines via scikit ^18,19^:

- “shLDA” refers to a Ledoit-Wolf shrinkage of the features before applying a Linear Discriminant Analysis (LDA) to improve the generalization performance of the classifier.
- ‘logisticReg” corresponds to a Logistic Regression classifier with a L2 penalty term.

### 2.5 Robustness assessment

We then conducted several tests to assess the robustness of the classification when randomly applying missing data (10% of the recording duration) to an increasing number of recordings. A total of 10 tests were performed, linearly increasing the number of subjects (4, 7, 10, 13, 16, 18, 21, 24, 27, 30) and the number of movements (18, 34, 51, 68, 85, 101, 118, 135, 151, 168) from which data points were removed. The removed data were subsequently filled using a moving median over a window of ten elements.

### 2.6 Statistical analysis

The classification accuracy was compared between the two features (i.e., kinectomes and PCA on time series). The accuracy values for the 30 movements obtained using the two different approaches were statistically compared using a permutation test based on 10,000 iterations. In each iteration, the values were randomly shuffled to create two new sets, and the absolute difference between the means of these two sets was calculated. Finally, the actual absolute difference in the mean accuracy between the two approaches was compared with the distribution of surrogate differences to obtain a statistical probability. Repeated measures ANOVA test was used to compare the accuracy performance in case of missing data. Kruskal-Wallis H-test was used to verify whether an effect of the classification method occurred. Multiple comparisons were corrected using False Discovery Rate (FDR) ^20^ with a significance level set at p < 0.05.

## 3. Results

### 3.1 Model prediction performance

The implemented method relied on the use of a Support Vector Machine (SVM) classifier. It aimed at distinguishing the 30 movements performed by 183 athletes. We compared the performance obtained by two different types of features: the kinectome (referred to as Kin-SVM), and the PCA applied on the time series (referred to as PCA-SVM). We observed that the classification accuracy of Kin-SVM was significantly larger as compared to the one obtained with PCA-SVM (mean ± standard deviation are 0.99 ± 0.01 and 0.96 ± 0.04 respectively, pFDR < 0.001). In particular, Kin-SVM presented a lower dispersion of the performance across the movements as compared to PCA-SVM (see Fig. 2). Such a tendency was confirmed by better performances measured via the area under the curve with AUCs of 1.00 for Kin-SVM. To get a more detailed description of the performances, we investigated the confusion matrices (see Fig. S1-S2), that show the classification accuracy for each movement and, in case of errors, indicate which movements were misclassified.

**Figure 2.**
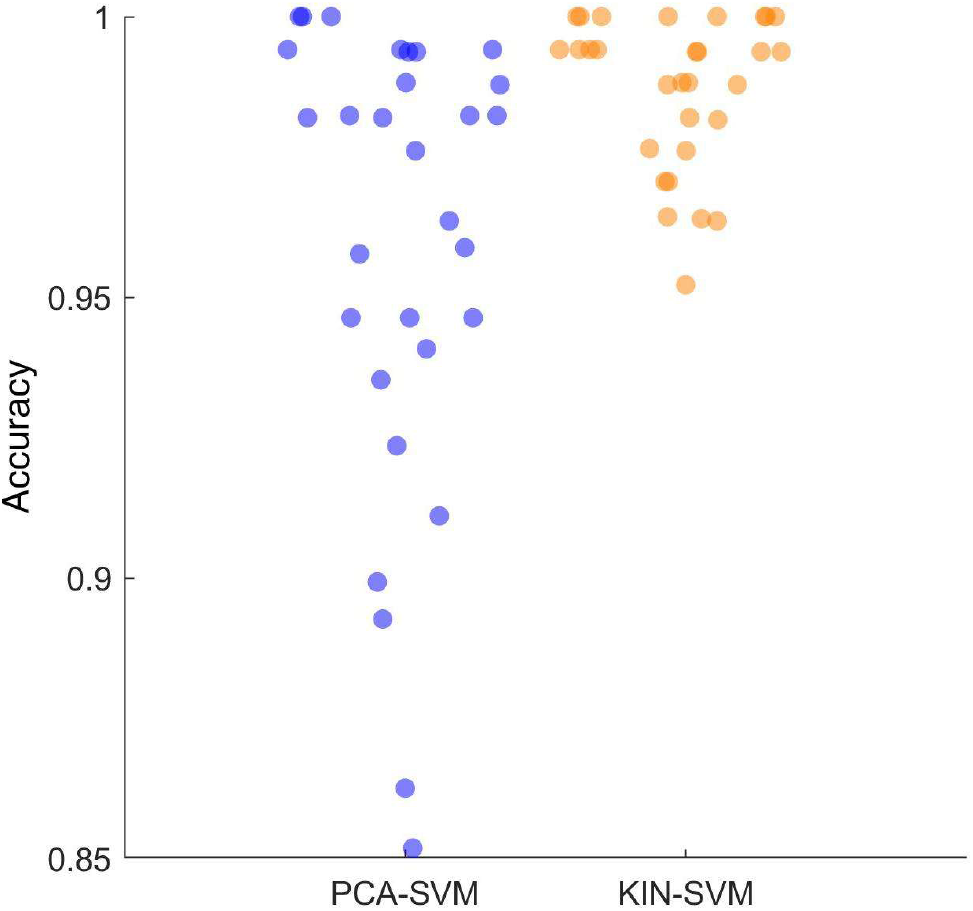
Classification accuracy comparison. Classification was performed via support vector machine (SVM), using features extracted using principal component analysis (PCA-SVM) and kinematic network analysis (KIN-SVM), separately. Each dot represents the accuracy score of each movement, averaged across 50 splits of cross-validation. KIN-SVM showed significantly higher classification accuracy than PCA-SVM (pFDR < 0.001).

Then, we replicated our results using two other classification algorithms (namely Linear Discriminate Analysis and Logistic Regression approaches) showing that they are resilient to this choice (see Fig. S3). We did not observe an effect of the classification method (Kruskal-Wallis H-test, p > 0.05).

### 3.2 Feature selected by the model

To get a better understanding of the obtained performances, we investigated the information that played the most important role in the classification by computing the features importance. In the case of PCA, the investigation of the feature importance is not straightforward. Indeed, even though we can have access to the actual components chosen across the splits, since the decomposition in principal component is performed for each split before the classification, these components result from a different decomposition and therefore cannot be compared. Therefore, we focused our study on the features relying on the kinectomes.

We observed a wide distribution of the importance of the features (Fig 3). This finding demonstrates that a subset of edges carries the information that is used to differentiate the different movements. To further investigate the information that played a major part in the classification, we focused on their formal identification and their visualization.

**Figure 3.**
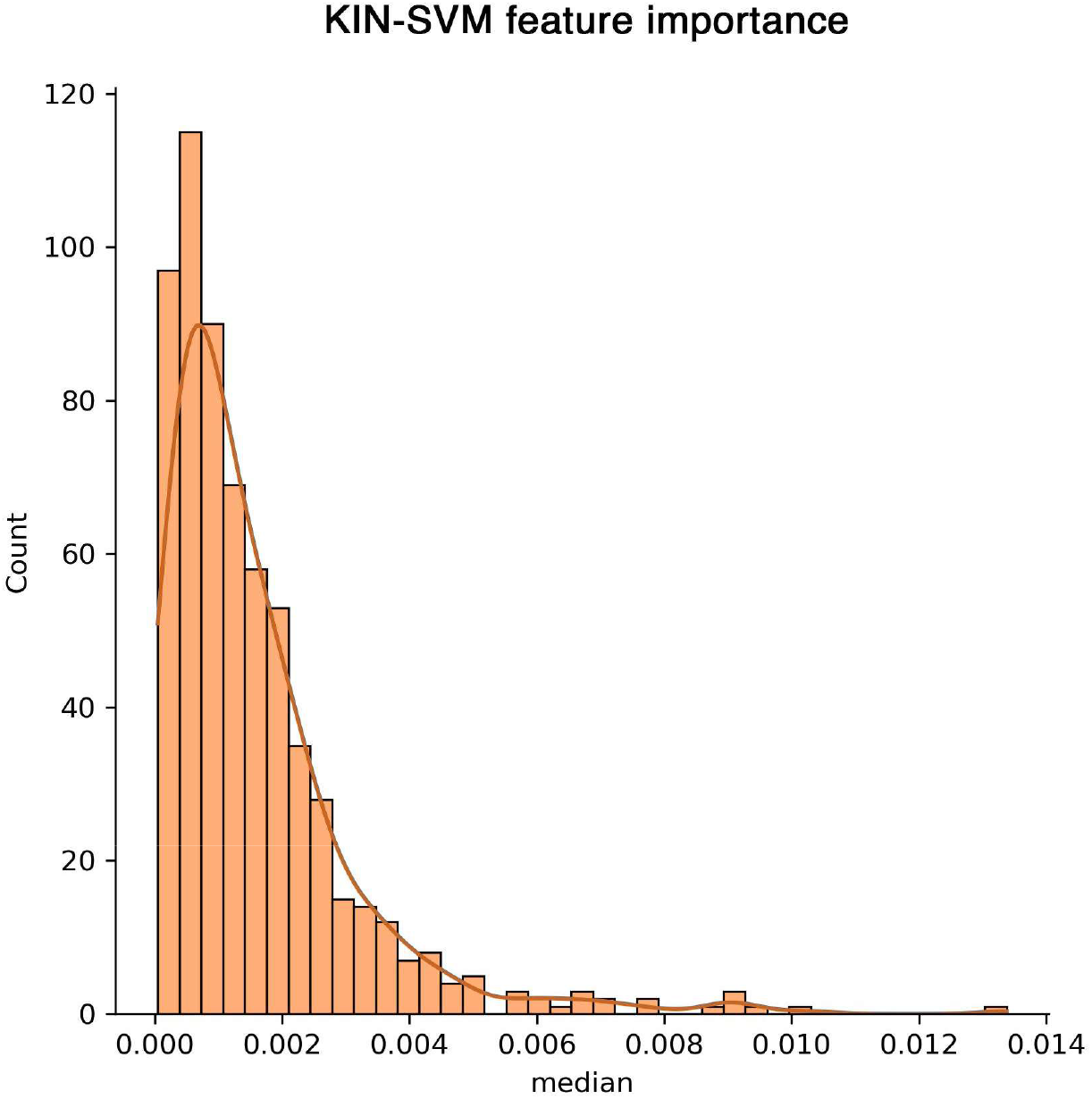
Feature importance distributions. The histograms represent the distribution of the feature importance values in support vector machine (SVM) classification, for features based on principal kinectomes (KIN).

Therefore, we focused our study on the visualization of the edges of the kinectomes that showed the higher feature importance rate. For this purpose, we used a circular plot that represents the interactions (i.e., the edges) between nodes, here the joints (see Fig. 4). The interactions with the highest importance involve mediolateral (ML) edges of the lower limbs and the trunk. Upper body joints are present too, but with lower values. Finally, the anteroposterior edges also prove highly important, immediately after the mediolateral ones.

**Figure 4.**
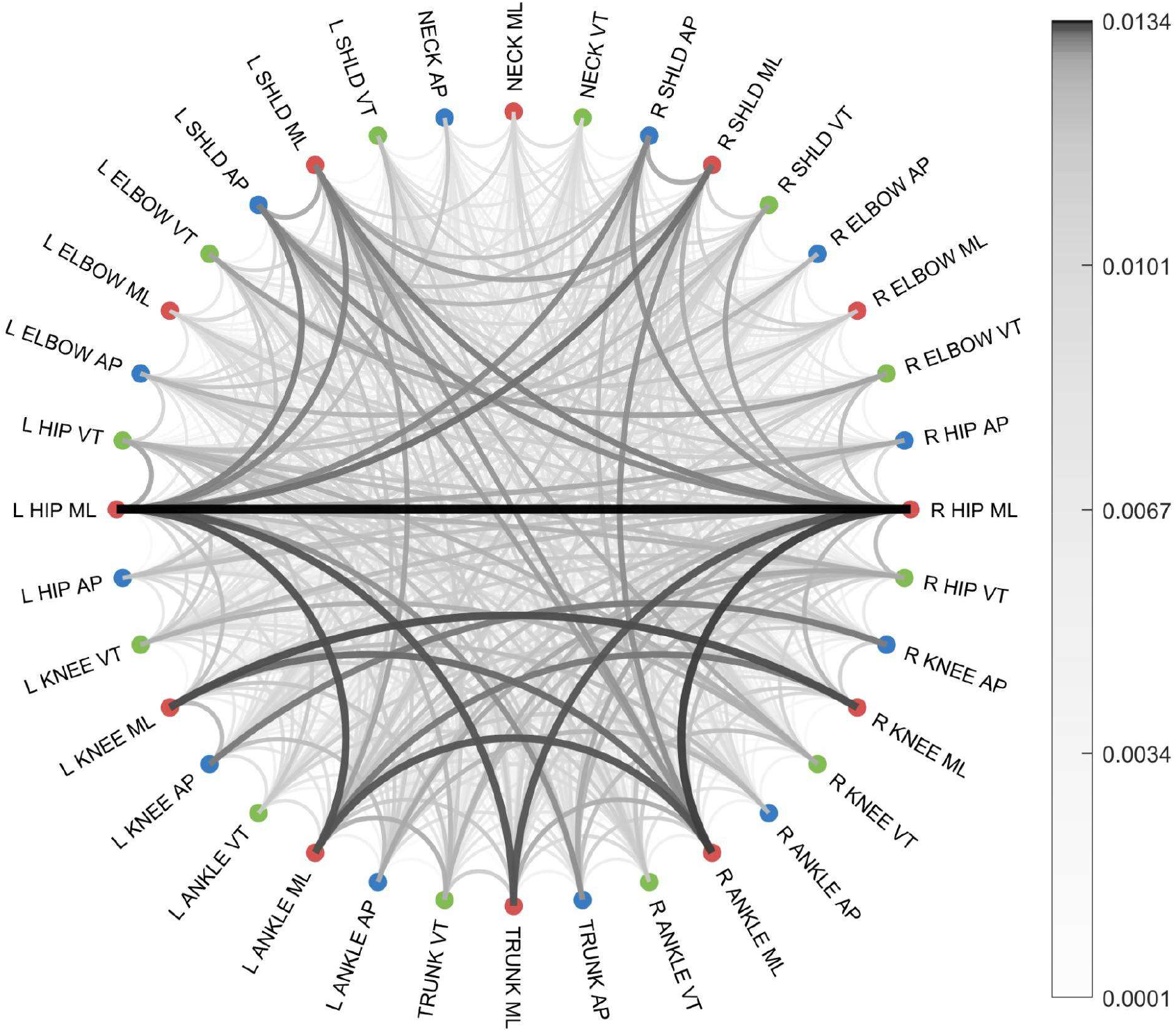
Network-based feature importance visualization. The circular plot offers a visual representation of the importance of the features used for classifying the 30 movements. Colored dots represent the node of the kinematic network (i.e., joints); movements on mediolateral, anteroposterior and vertical axes were represented by red, blue, and green colors. The links between nodes represent the edges of the network that, in turn, represent the coordination between two given joints. The higher the importance of a movement for classification, the darker the color and the thicker the line.

### 3.3 Robustness

We further examined how the presence of missing data impacts classification performance to assess the robustness of the features under these conditions. The results in Figure 5 show that PCA-SVM performs worse compared to KIN-SVM (F(9,522) = 49.69, p < 0.001). Additionally, the performance of the PCA-SVM decreases as the amount of missing data increases (F(9, 261) = 51.49, p < 0.001), whereas the Kinectome appears unaffected in all cases (F(9, 261) = 1.07, p = 0.382).

**Figure 5.**
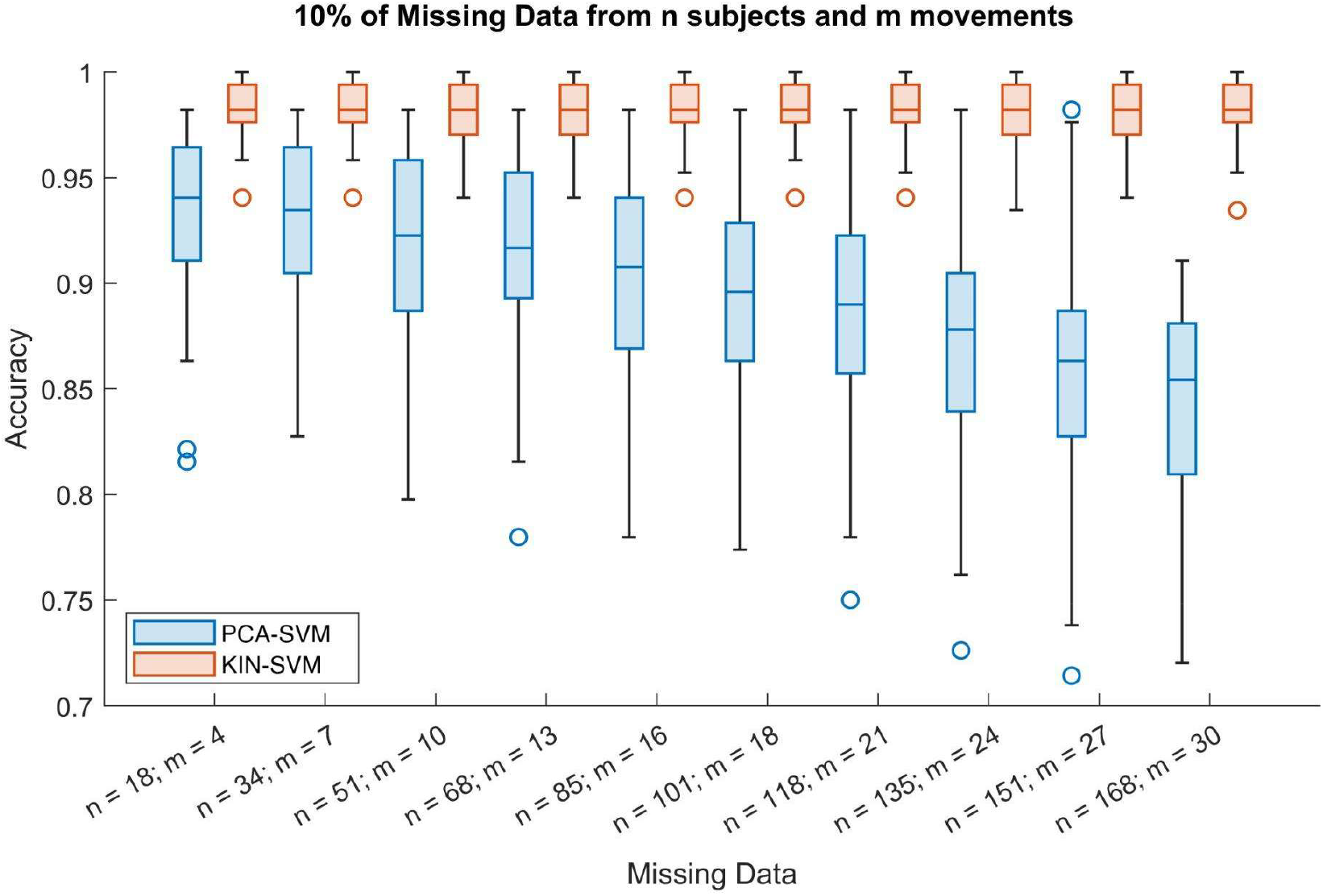
Feature resilience to time series with missing data. The box plots show the accuracy in classifying 30 different movements using a support vector machine (SVM) while randomly removing increasingly larger amounts of data ten times. Specifically, with each repetition, more data was removed from a larger number of participants (i.e., as shown on the x-axis). The data was removed from the original time series, while classification was performed on features derived from principal component analysis (PCA) and kinectomes (KIN).

### 3.4 First and second derivatives

We also evaluated the performance when the features were calculated from the derivatives of the joint excursion time series. In movement analysis, derivative measures such as velocity and acceleration are often used, as they provide complementary information useful for data investigation. The results shown in Figure 6 indicate that the accuracy performance drops significantly with the simple PCA on time series, and worsens as the derivative order increases (velocity: 0.93 ± 0.19; acceleration: 0.11 ± 0.16). In contrast, with the kinectome, the result remains mostly stable, with only a slight decrease when applied to the higher-order derivative (velocity: 0.99 ± 0.01; acceleration: 0.98 ± 0.04).

**Figure 6.**
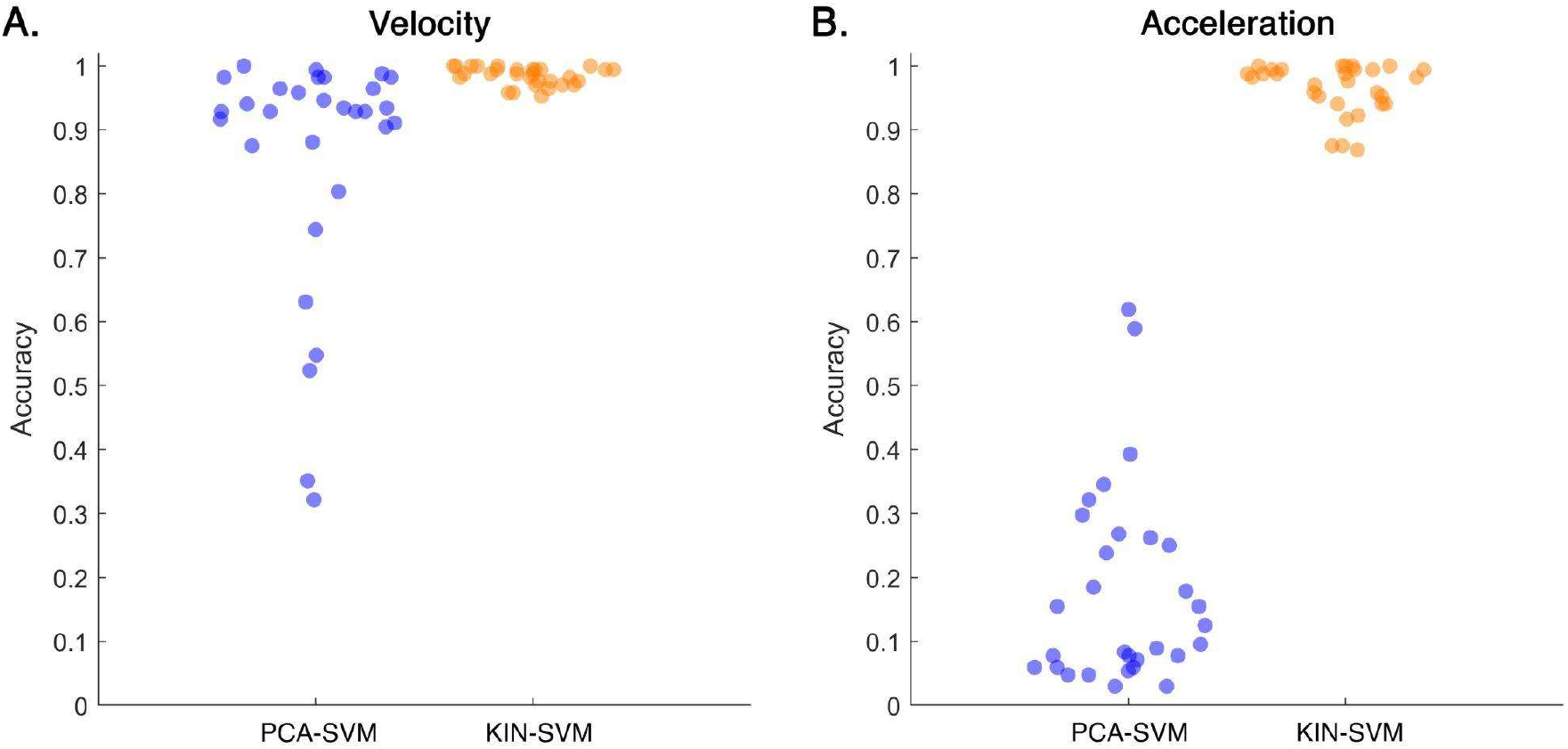
Classification accuracy with derivative data. Classification was performed via support vector machine (SVM), using features extracted using principal component analysis (PCA) and kinematic network analysis (KIN), starting from the first (velocity) and the second (acceleration) derivative of the time series. Each dot represents the accuracy score of each movement, averaged across 50 splits of cross-validation. KIN-SVM showed significantly higher classification accuracy than PCA-SVM (pFDR < 0.001).

## 4. Discussion

In this study, we tested the classification of different movements utilizing the kinectome. This novel technique conceptualizes the human body as a network, where joints are the nodes and the coordination between pairs of joints is represented by edges. The classification performance of the kinectome for different movements was benchmarked against PCA as a dimensionality reduction method. The tested dataset consisted of time series of joint excursions from multiple human body joints during the execution of 30 movements performed by 168 individuals. Our main hypothesis is that network-based features, in addition to the immediacy in visual representation, and thus in interpretation, would lead to better classification performance compared to PCA-based features. The results confirm our hypothesis, showing better and more robust classification performance.

The literature reports relatively high classification performance for movements, often ranging between 70% and 95% accuracy ^21^. Hence, we expected a good performance for the classification based on PCA. Nevertheless, PCA showed greater variance in accuracy when considering all movements (i.e., some movements had an accuracy below 90%). In contrast, the kinectome displayed a higher average accuracy with a reduced dispersion compared to PCA (standard deviation ±0.01 and ±0.04, respectively), never falling below 90%, regardless of the movement. This suggests stability in the performance of kinectomes’ features even in relation to the variety of movements observed, although still limited to the 30 movements in this dataset. Comparing different studies can be challenging, as the observed movements, features, classification algorithms, and other variables are often not comparable across studies. For instance, Gonzalez et al., reported that studies on the recognition of human activities show accuracies ranging from 70 to 95% ^21^. Similarly, a classification analysis aimed to distinguish standing, walking, and running, in 5 subjects, based on acceleration data from inertial sensors, electromyography, and force plate data, reached 86.4% classification accuracy. This performance was achieved by using PCA to reduce data dimensionality and SVM to classify the movements ^21^. Zebin et al., using a dataset consisting of 6 actions performed by 12 individuals, carried out various classification tests based on five inertial sensors placed on the lower limbs ^22^. They compared the performance of numerous feature extraction approaches and classification methods (SVM, multilayer perceptron, and convolutional neural networks), obtaining classification accuracies of up to 97%, with SVM showing an accuracy equal to 96.4%. A similar accuracy was reported by Mascret et al., in a study including eight movements performed by four participants ^2^. The authors classified the movements with a 97.7% accuracy using an SVM classifier, utilizing time series from 3D accelerometers, gyroscopes, and magnetometers, placed on bust, thigh, and tibia. A common limitation of studies similar to ours is the evaluation of a limited set of movements, typically ranging from four to eight, performed on a small sample of participants, usually between four and twelve. In our study, leveraging the dataset published by Zhao et al. ^15^, we tested our approach on 168 subjects performing 30 different movements, achieving the best classification performance (i.e., accuracy = 99%) when using whole-body coordination values as features, obtained through the network theory approach. We replicated these findings with two more classification algorithms, namely LDA and Logistic Regression (Fig. S3). Here, our approach was focused on the use of inertial information from time series. Given the performance obtained with the first and second derivatives, a future work would consist in the enrichment of the information used as features through more sophisticated classification tools, similarly to ^23^

Furthermore, our approach emphasizes the interpretability of the features. Understanding which elements contribute to classification can provide valuable insights for movement assessment, which can be useful in several contexts (e.g., sports and clinical fields) ^24,25^. The interpretability of motor features depends on the type of features selected. Typically, the data features stem from dimensionality reduction algorithms (such as PCA) or the combination of very different parameters ^26^. In the first case, the interpretation proves challenging, as the low dimensional variables correspond to a combination of the original variables. The second case includes studies that put together very different variables, use movement-specific variables, or consider only certain body parts without accounting for their interdependencies (i.e., the coordination) that are a prerequisite to generate the correct movement ^27–29^. This might bring a lack of consistency in the results, and the outcomes may depend on the particular features taken into account. With our network theory-based approach, we obtained a feature that is not movement-specific, include information on coordination ^30^, and offer an easy-to-interpret visualization of the results, which is suitable for non-specialized users. Our feature importance analysis highlights the edges of the coordination matrices that are most informative for classification, pointing to specific joints along specific planes of motion (Fig. 4). This approach provides a readily usable tool, which might also be optimized for online and real-time investigations to recognize movements but also to evaluate their role from a biomechanical perspective.

Moreover, our approach demonstrated high reliability even in case of missing data. Although it is common to compensate for data loss using methods such as interpolation, in an online environment this would require an alternative approach. With the kinectomes, we observed classification performances that are robust to data loss. Furthermore, when analyzing movements, relevant information is contained in the derivatives of data (e.g., velocity and acceleration), and our analysis showed that the kinectomes performed better than PCA even in this case as well.

Several limitations of our work should be highlighted. Firstly, it would be essential to demonstrate high cross-dataset accuracy to generalize the classification across different datasets and motion capture systems. Equally important, our approach should be tested in real-time environments. Future applications, in addition to addressing these limitations, could also consider clinical settings to evaluate the effectiveness of this approach in classifying and assessing movement execution in clinical populations for supporting diagnosis ^31,32^. Furthermore, machine learning techniques based on network-derived features are widely employed in neuroscience, particularly in motor imagery research ^33^. Future studies could integrate these methods to enhance the classification of both approaches by leveraging data from the imagined and executed components of movement. Additional applications in sports may include integrating physiological parameters and performance staging to differentiate professional athletes from amateurs ^34^.

In conclusion, in this study, we investigated the accuracy of alternative features for movement classification. As a first proof-of-concept, we focused our analysis on the key criteria to meet (namely, classification performance, explainability, and robustness to missing data) with the simplest classification algorithm, i.e. the support vector machine. Our study has found that utilizing features based on network analysis offers numerous advantages. Firstly, the classification accuracy is superior to other traditionally used methods in many regards. The accuracy of features based on the kinectome is higher regardless of the classifier, it does not drop when using kinectomes based on derivatives, and is resilient to data loss. Additionally, the kinectome allows for the visualization of features and their importance, providing topographical interpretations concerning the anatomy and kinematics of the human body. Overall, these findings could pave the way to automated movement recognition and evaluation tools, aiming to support sports coaches on the field, physicians and rehabilitators in everyday activities, or any other workers concerned with human motion recognition.

## Acknowledgments

This work was supported by:

University of Naples Parthenope, Via Ammiraglio Ferdinando Acton, 38, 80133, Napoli (Italy). P.IVA: 01877320638 - C.F. 80018240632. Governo Italiano Ministero per lo sviluppo Economico, ACCORDI PER INNOVAZIONE. Approccio User-friendly integrato per Diagnosi, Assistenza e Cura Efficaci—AUDACE. CUP: B69J23006050007.

European Union “NextGenerationEU”, (Investimento 3.1.M4. C2), project IR0000011, EBRAINS-Italy of PNRR.

## Declaration of interest

The authors declare they have no competing interests.

## Author contribution statement

ETL conceptualized the study, led the investigation, performed the main analyses, and drafted the manuscript. RM assisted in drafting the manuscript and provided theoretical insight into movement analysis. MDL managed data curation, supported the analyses, created the figures, and revised the manuscript. FT was responsible for data curation and reviewed the manuscript. GS reviewed the manuscript and secured the funding, while PS contributed to software development for data management and analysis, participated in the investigation, and revised the manuscript. MCC led the investigation, performed the main analyses, drafted the manuscript, supervised the work and contributed statistical expertise. All authors have read and approved the final version of the manuscript and agree with the order of presentation of the authors.

## Data statement

The data used in this manuscript are provided by Zhao et al., 2023, and are publicly available at: https://doi.org/10.1038/s41597-023-02082-6

## Notes

https://doi.org/10.1038/s41597-023-02082-6

### Competing Interest Statement

The authors have declared no competing interest.

