## Supplementary material for "Artificial Intelligence for automatic movement recognition: a network-based approach": Fig. S

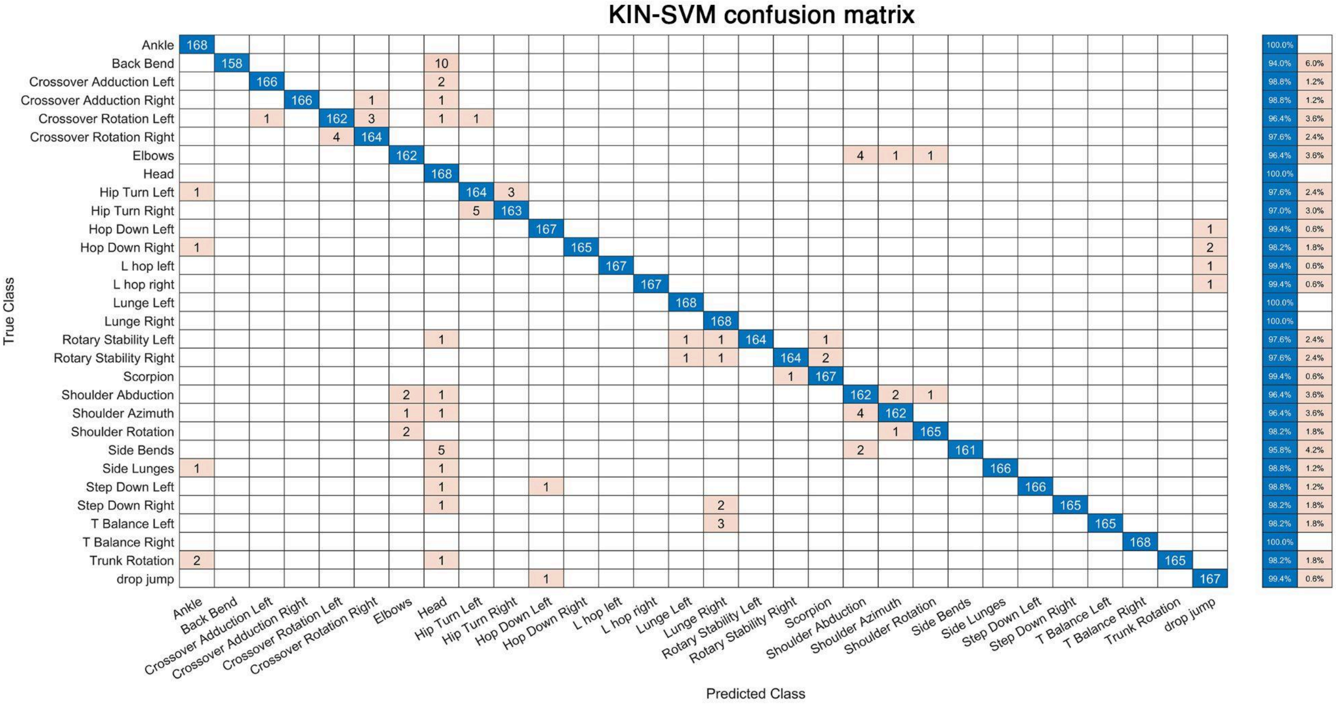


Figure S1. Confusion matrix of support vector machine classification based on kinectomes.

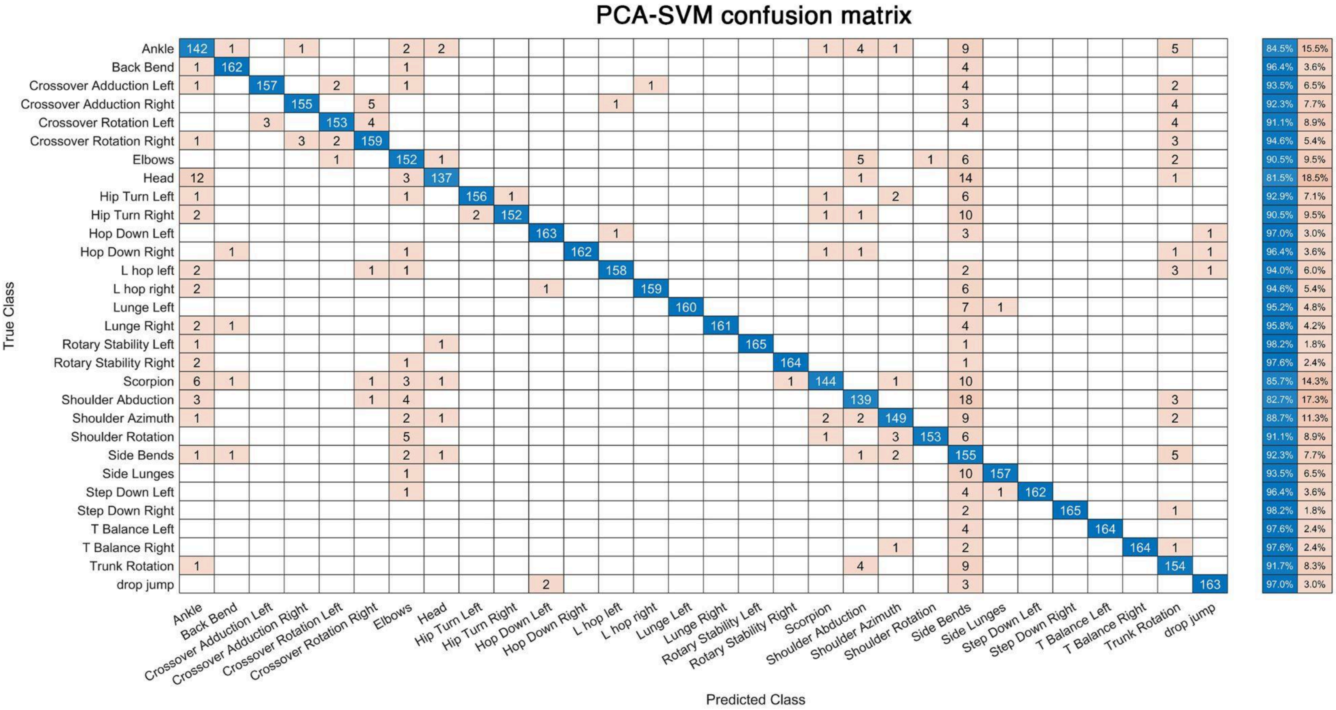


Figure S2. Confusion matrix of support vector machine classification based on principal component analysis (PCA).

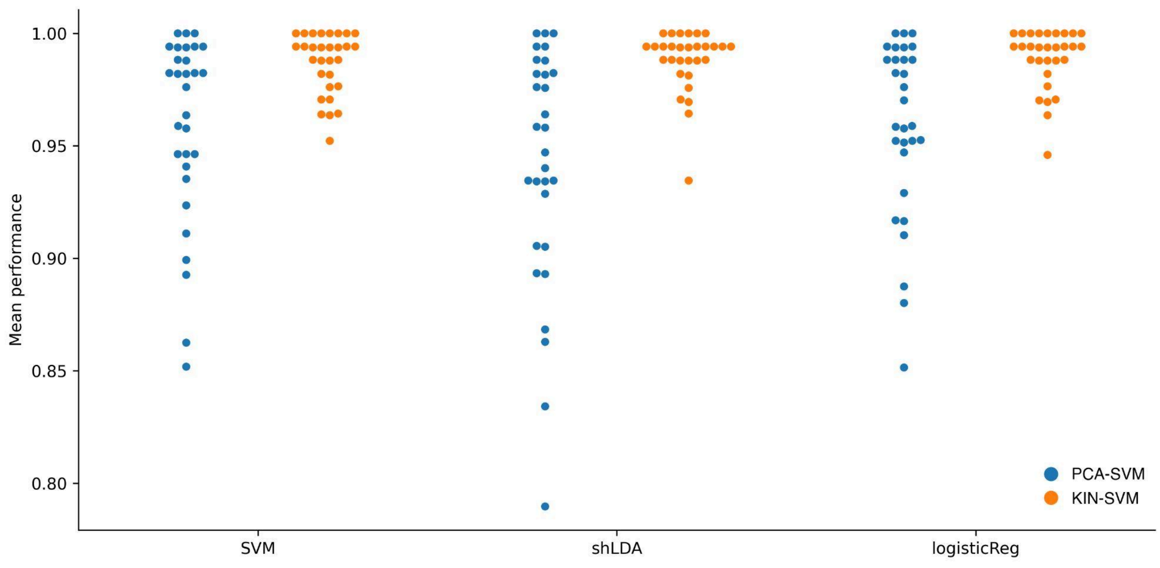


Figure S3. Classification accuracy with different classification algorithms.
